# Impact of *Candida auris* infection in a neutropenic murine model

**DOI:** 10.1101/731281

**Authors:** Steven R. Torres, Amber Pichowicz, Fernando Torres-Velez, Renjie Song, Navjot Singh, Erica Lasek Nesselquist, Magdia De Jesus

## Abstract

*Candida auris* has become a global public health threat due to its multidrug resistance and persistence in hospital and nursing home settings. Although a skin colonizer, *C. auris* can cause fatal bloodstream infections and most patients succumb to multi-organ failure. Currently, there are limited animal models to study the progression of *C. auris* infection. Here we compare two murine models of neutrophil depletion using monoclonal antibodies 1A8, anti-Ly6G^+^ and RB6-8C5 anti-Ly6G^+^-Ly6C^+^. We also compare inoculums of 10^7^ and 10^8^ as well as the intravenous and gavage routes of infection. The results reveal that neutrophil depletion in BALB/c mice is sustained long-term with the 1A8 antibody and short-term with RB6-8C5. Target organs were kidney, heart and brain as these had the highest organ fungal burden in neutrophil depleted and to some extent in infected control mice. We found that *C. auris* is shed in urine and feces for neutrophil depleted mice. The gavage model is not an ideal route as dissemination was not detected. Eight days post *C. auris* infection all surviving mice display a unique behavioral phenotype characterized by torticollis and tail spin that progresses to head bobbing and body curling like phenotype by day 22, that continues to persist even 104 days post infection. Lastly, we found *C. auris* remains in tissues of infected control mice, 34 plus days post infection suggesting that *C. auris* stays present in the host without causing disease but becomes opportunistic upon a change in the hosts immune status such as neutrophil depletion.

## Introduction

Within a decade, the fungal organism *Candida auris* has become a major public health concern because of its unpredictable multidrug resistance (MDR) phenotype, persistence in hospital and nursing home settings, transmissibility and high mortality (1–5). Although its origin is currently unknown, its rapid independent and simultaneous worldwide emergence has given rise to four distinctive clades (1–5). *C. auris* is resistant to an already limited arsenal of antifungal classes that includes the azoles and polyenes. Most recently, a collection of 54 isolates obtained by the Centers for Disease Control (CDC) from India, Pakistan, South Africa, and Venezuela revealed that 93% of isolates were resistant to fluconazole, 35% were resistant to amphotericin B, and 7% were pan resistant to the echinocandins, a third class of antifungals that are currently used to treat *C. auris* infections (6, 7). *C. auris* is a persistent skin colonizer and can last on surfaces for up to a month without proper disinfection methods (8, 9). Since its identification, progress has been made in the areas of epidemiology, the development of rapid and specific diagnostics such MALDI TOF and real-time PCR, and in determining appropriate disinfection methods (1, 10–12).

Although *C. auris* can colonize all individuals, the risk factors are similar to other types of *Candida* infections that include individuals who have compromised immunity such as those with cancer, diabetes, transplants as well as a history of abdominal surgery, presence of central venous catheters, ventilators and prolonged antibiotic treatment (1, 13). Case studies have highlighted that patients who ultimately succumbed from systemic *C. auris* infections exhibited renal failure, heart failure with endocarditis and ventriculitis respiratory arrest, pneumonia and multi-organ failure (14–17). In two case studies, 72 and 74- year old patients were admitted with alteration of mental status and unsteady gait. However, there was no clear indication whether this was due to *C. auris* infection (14, 18). Currently, there are *Drosophila melanogaster* and *Galleria mellonella* models for *C. auris* infection. There are also a limited number of murine models (19, 20). The current murine models for *C. auris* infection have used immunocompetent and immunosuppressed models. In the immunocompetent models, mice are infected with inoculums that range from 10^5^ to 10^7^ cells per animal (21, 22). However, there appears to be a difference in survival between the studies by Fahkim et al. and Wang et al. (21, 22). Fakhim et al. used ICR an outbred murine strain and infected them using two clinical isolates from the South Asian clade, that were characterized as non-aggregative and virulent (21). Fakhim et al. reported a 60%-70% mortality rate by day 17 post-infection (21). Wang et al. used BALB/c an inbred strain and infected with isolates also from the South Asian clade (22). These mice did not succumb to infection after 14 days post-infection. Wang et al. attributes this discrepancy to the difference in mouse strains and differences in *C. auris* isolates (22). The murine models of immunosuppression use the chemotherapeutic agent cyclophosphamide, or a combination of cyclophosphamide and cortisone acetate (23, 24). These models work well in achieving infection and for evaluating the efficacy of a particular treatment such as antifungal and vaccine candidates against *C. auris* (23, 24). However, a limitation of using cyclophosphamide is that it depletes cells of both the innate and the adaptive immune system (25). At high doses of cyclophosphamide ≥200mg/kg, these deplete Tregs, CD8^+^ DCs, B-cells and CD4^+^ and CD8^+^ T-cells that may be important in immunological studies in response to *C. auris* (23, 24, 26–29).

To develop a model of *C. auris* infection as a tool to understand pathogenesis, we tested several variables in BALB/c mice such as whether we could obtain infection by a sustained neutrophil depletion with anti-neutrophil antibodies 1A8 and RB6-8C5. We also tested different inoculums of 10^7^ and 10^8^ *C. auris* cells and different routes of infection such as intravenous and gavage. Here we report that in addition to finding differences in the tested variables we also found a behavioral phenotype that becomes evident in mice 8-13 days post i.v. infection with *C. auris* regardless of immune status that does not occur in control mice. Lastly, we found *C. auris* remains in tissues of infected control mice 34 plus days post infection suggesting that *C. auris* stays present in the host without clinical disease and may become opportunistic upon a change in immune status. Moreover, detection of yeast clusters by IHC in the renal pelvis of these chronic carriers indicates that *C. auris* is actively being shed in the urine weeks after the clinical course.

## Results

We depleted neutrophils using two distinct neutrophil depleting antibodies: 1)1A8 an anti-Ly6G^+^ antibody that only depletes neutrophils and 2) RB6-8C5 an anti- Ly6G^+^-Ly6C^+^ that depletes Ly6G^+^and Ly6C^+^ neutrophils, dendritic cells, and subpopulations of lymphocytes and monocytes (30). We depleted neutrophils by administering 1A8 or RB6-8C5 antibodies at 24 hours prior to *C. auris* infection and every 48 hours thereafter (31). To measure the degree of neutrophil depletion and its sustainability, we used flow cytometry and analyzed blood and spleen that represent circulating and tissue resident neutrophils. We also collected urine and fecal samples to test for the presence of *C. auris* throughout the course of infection (Fig. 1A). At days 2,4, and 6 in spleen both antibodies worked well in depleting Ly6G^+^-Ly6C^+^ double positive neutrophils to 0.20%, 0.29%, and 0.27% respectively for 1A8, and 0.11%, 0.08%, and 0.37% respectively for RB6-8C5. However, by day 39, 1A8 continued to deplete Ly6G^+^-Ly6C^+^ double positive neutrophils to 0.51% whereas RB6-8C5 did not and Ly6G^+^-Ly6C^+^ double positive neutrophils increased to 7.66% despite that the antibody was administered every two days (Fig. 1B). We also observed a similar trend at days 2,4, and 6 in blood with both antibodies working well in depleting Ly6G^+^-Ly6C^+^ double positive neutrophils to 0.01%, 0.11%, and 0.03% respectively for 1A8, and 0.02%, 0.05%, 0.04% respectively for RB6-8C5 (Fig.1C). By day 39, 1A8 continued to deplete Ly6G-Ly6C double positive neutrophils but RB6-8C5 did not with Ly6G-Ly6C double positive neutrophils going up to 11.56% (Fig. 1B and C). With the understanding that mice eventually compensated the Ly6C^+^-Ly6G^+^ cells with the use of RB6-8C5 antibody, we chose to compare the progression of *C. auris* infection in both the 1A8 and RB6-8C5 neutrophil depleted models. As 1A8 represents long term depletion while RB6-8C5 represents short term depletion.

**Figure 1.**
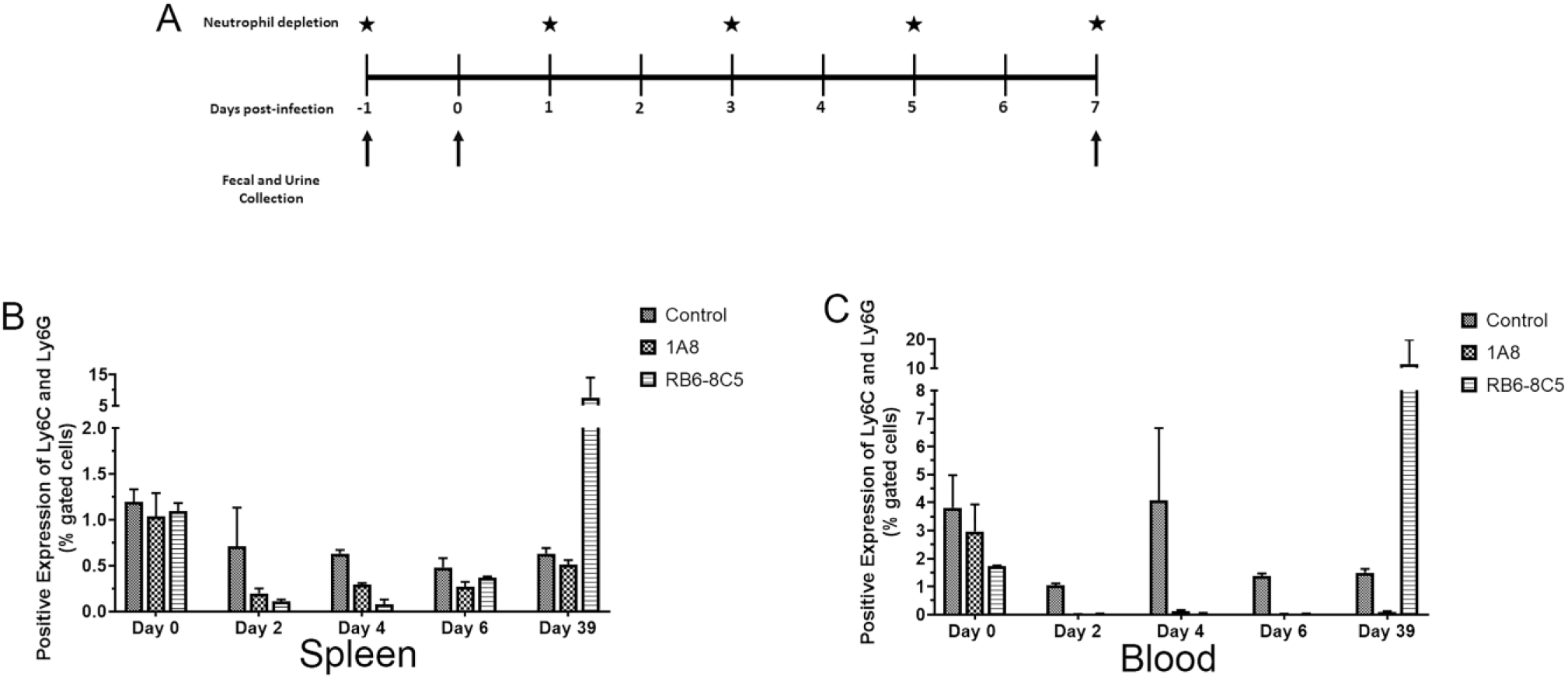
Depletion of neutrophils using monoclonal antibodies 1A8 and RB6-8C5 shows that 1A8 is optimal. (A) Representative strategy of neutrophil depletion. BALB/c mice were injected with 1A8 or RB6-8C5 at day −1 and day 0. At day 1, mice are infected with 10^7^ or 10^8^ *C. auris* by i.v. or gavage. Neutrophil depletion antibodies were then administered every 2 days. Fecal and urine samples were collected on days −1, 0 and 7. (B) Comparison of the efficacy of neutrophil depletion between 1A8 and RB6-8C5 in spleen, depletion of Ly6G^+^-Ly6C^+^ double positive were assessed. (C) Blood shows a similar trend as the spleen suggesting that 1A8 is optimal to sustain neutrophil depletion.

Survival studies in *C. auris* i.v. infected mice showed that neutrophil depletion with either 1A8 or RB6-8C5 at a dose of 10^8^ was fulminant to all mice by days 3 and 4 (Fig. 2A). We noted that RB6-8C5 treated mice infected with an inoculum of 10^8^ by day 1 displayed a hunched posture, swollen shut eyes and rough coat as previously described (21), (Fig. 2B). By day 1 post infection the 1A8 treated mice, with a 10^8^ inoculum, showed a normal posture and slightly rough coat. However, both neutrophil depleted groups eventually died by days 3 and 4 and therefore did not represent an ideal model to study the course of infection (Fig. 2A). Mice infected via gavage did not show signs of illness and there was 100% survival regardless of which neutrophil depletion antibody or inoculums that were used (Fig. 2A). At a dose of 10^7^, we noted differences in the appearance of mice, depending on the neutrophil depletion antibody that was used. The RB6-8C5 treated mice showed signs of illness as early as day 1 as determined by a rough erect coat, hunched-back and had lost 20% loss of their weight by day 7 (Fig. 2F). At this inoculum, using RB6-8C5, there was a 60% survival rate and the surviving mice recovered 15% of their starting weight by day 16 which may be due to the immune system overcompensating for the neutrophil depletion. The mice treated with 1A8 showed signs of illness by days 5 and 6 determined by a rough erect coat, hunched-back and 15% loss of their weight by day 7. For 1A8, 40% of the mice survived, and those recovered 5% of their weight by day 16 (Fig. 2C and 2E). Infected control mice at inoculums of 10^8^ and 10^7^ showed slightly rough coats by days 1 and 2 but there was 100% survival in both groups. At the 10^8^ inoculum mice lost 3% body weight by day 10 and the mice inoculated 10^7^ lost 5% weight by day 17 (Fig. 2A and 2E).

**Figure 2.**
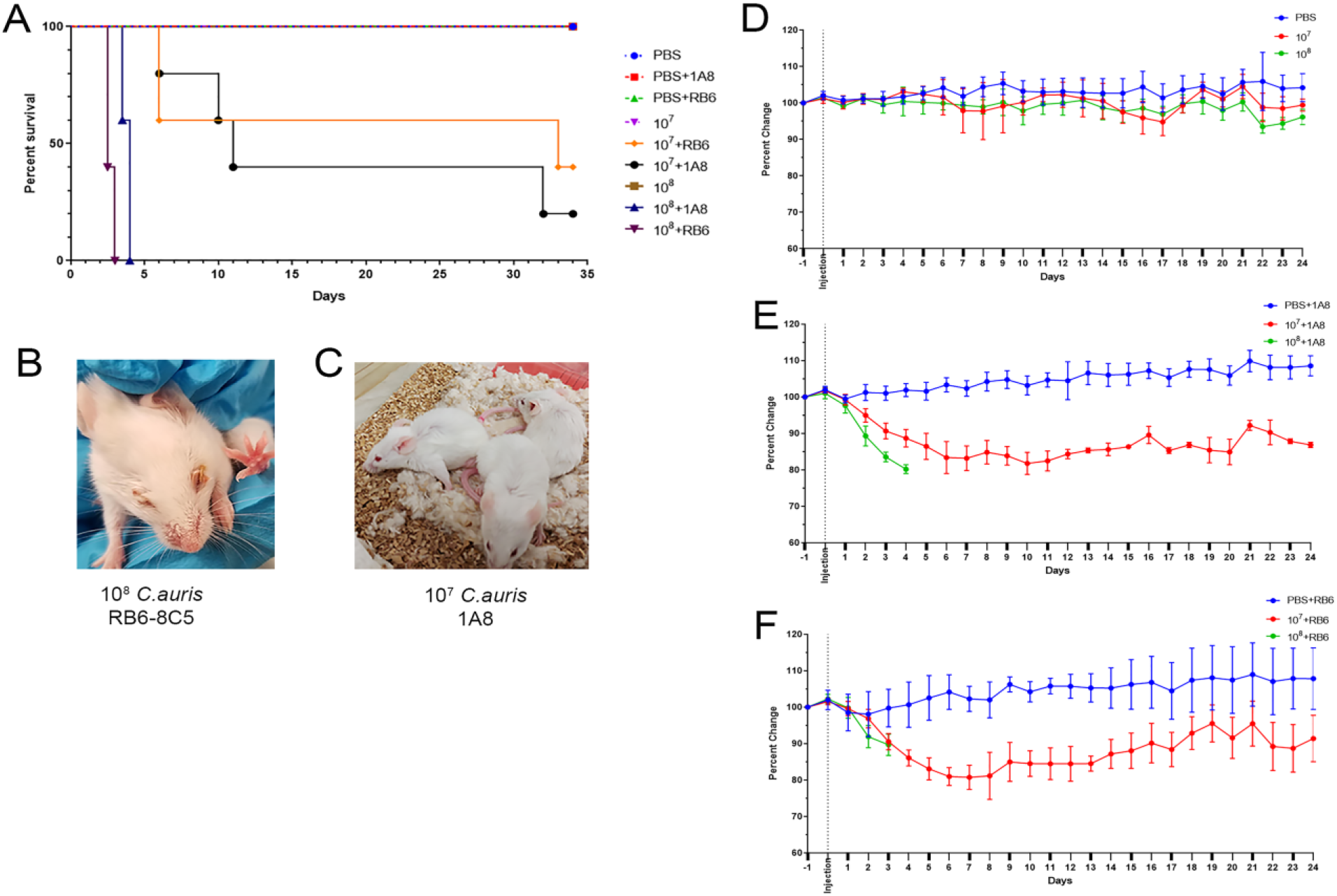
Survival study comparing immunocompetent mice and neutrophil-depleted mice infected with *C. auris*. (A) By day 34, PBS injected mice as well as immunocompetent mice infected with 10^7^ or 10^8^ *C. auris* cells had 100% survival. All neutropenic mice infected with 10^8^ succumbed to infection by day 3 for RB6-8C5 and day 4 for 1A8 treated mice. RB6-8C5 infected mice infected with 10^7^ *C. auris* had a survival rate of 40% while 1A8 mice infected 10^7^ *C. auris* had a survival rate of 20%. n=5 per group (B) Representative RB6-8C5 neutrophil depleted mouse infected with 10^8^ *C. auris* had swollen shut eyes (C) Representative 1A8 neutrophil depleted mouse infected with 10^7^ *C. auris* showed hunched posture and rough coat. (D-F) Mice in the **s**urvival studies were weighted for 24 days. All immunocompetent mice had similar weights throughout the course of the experiment. 1A8 infected with 10^7^ had an 15%-18% weight loss while RB6-8C5 infected with 10^7^ had 20% weight loss. By day 13, RB6-8C5 treated mice infected with 10^7^ begin to recover their weight by 10%-12%.

We measured fungal organ burdens 7 days post infection in mice that were infected with an inoculum of 10^7^ *C. auris* cells to determine which organs were affected by *C. auris*. We also compared differences based on the neutrophil depletion antibody that was used and the route of infection. For i.v. infected mice, we found that the kidney, heart, and brain were the organs with the highest fungal burden regardless of which neutrophil depleting antibody was used. For kidney and heart in i.v. infected mice, we found a fungal organ burden of 10^5^ CFU/g and 10^4^ CFU/g in the brain (Fig. 3). In other organs such as the bladder, uterus, spleen, cecum, small intestine, stomach and lungs from i.v. infected mice, we found modest levels of fungal burden that ranged from 10^1^-10^3^ CFU/g. We found enhanced organ fungal burden in neutrophil depleted mice with RB6-8C5 (Fig. S1). We did not find organ fungal burden in either kidney or brain in the gavage model but we found modest fungal burdens in the cecum, small intestine and stomach (Fig. S1). To confirm that the colonies that were detected on our plates were *C. auris*, we performed MALDI-TOF on samples from harvested organs. The results reveal that all tissues were infected with *C. auris* (Table S1).

**Figure 3.**
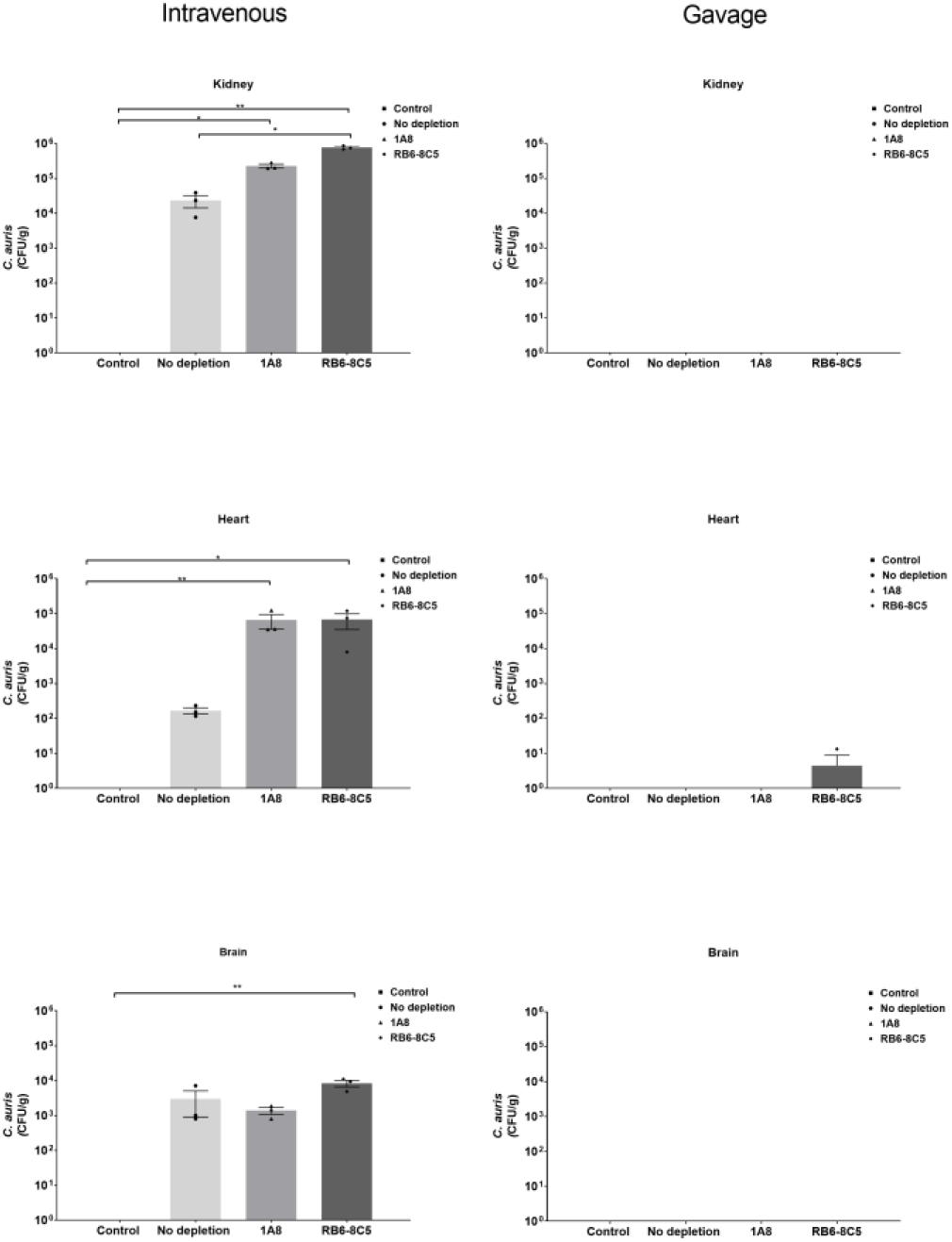
*C. auris* targets kidney, heart and brain by i.v. route of infection. CFU/g of tissues 7 days post-infection from mice infected by i.v. and gavage with 10^7^ *C. auris.* The major target organs in both neutrophil depleted and immunocompetent mice infected by the i.v. route were with CFU/g ranging from 10^4^-10^6^. The oral gavage route is not optimal to establish a *C. auris* infection in these organs. N=5 mice per group. A nonparametric one-way analysis of variance (ANOVA) with Kruskal-Wallis test followed by uncorrected Dunn’s multiple comparison test was done using ns P > 0.05, *P ≤ 0.05, **P ≤ 0.01, ***P ≤ 0.001, ****P ≤ 0.0001

In order to evaluate and score *C. auris* infected tissues, we determined that the fixation time frame in 10% formalin was 24 hours to fully inactivate *C. auris*. We determined this by culture and by counting CFU in infected formalin fixed tissues (Fig. S2). Histology was done using the standard hematoxylin-eosin (H&E) staining as well as by immunohistochemistry (IHC) staining for *C. auris* with an anti-*Candida albicans* polyclonal antibody that we found to cross react (Thermofisher, Waltham MA). Significant histopathological changes were observed in 1A8 and RB6-8C5 treated neutrophil depleted mice that were challenged with *C. auris* intravenously. We also found moderate changes in i.v. infected control mice. For all three groups, the kidney, heart, and brain were most affected, followed by the liver, gastrointestinal tract, and spleen (Fig 4 and Table 1). In the kidneys, there was a multi-focal-to-coalescing necrotizing tubulointerstitial nephritis, often involving the renal pelvis and capsule. The inflammatory component consisted mostly of mature and immature (band cells) neutrophils and macrophages, with prominent intralesional yeast. These lesions were significantly more severe and extensive in 1A8 and RB6-8C5 treated mice in comparison to the infected control mice. In these infected control mice the lesions consisted mostly of scattered microabscesses and focally extensive areas of inflammation. Similarly, changes in the heart were global and more prominent in infected 1A8 and RB6-8C5 treated neutrophil depleted mice. These changes consisted of multi-focal-to-coalescing necrotizing myocarditis, predominantly infiltrated by macrophages and scattered neutrophils. This infiltrate was often expanding between myofibers in areas where the organ architecture was still intact. In the infected control mice, the severity of inflammation was minimal and necrosis was absent. Brain damage was more prominent in infected animals treated with the 1A8 antibody. In these animals, there were multiple microabscesses and large foci of necrosis and edema throughout the brain. These areas of inflammation were infiltrated by macrophages, glial cells, and a few neutrophils (Fig 4 and Table 1).

**Figure 4.**
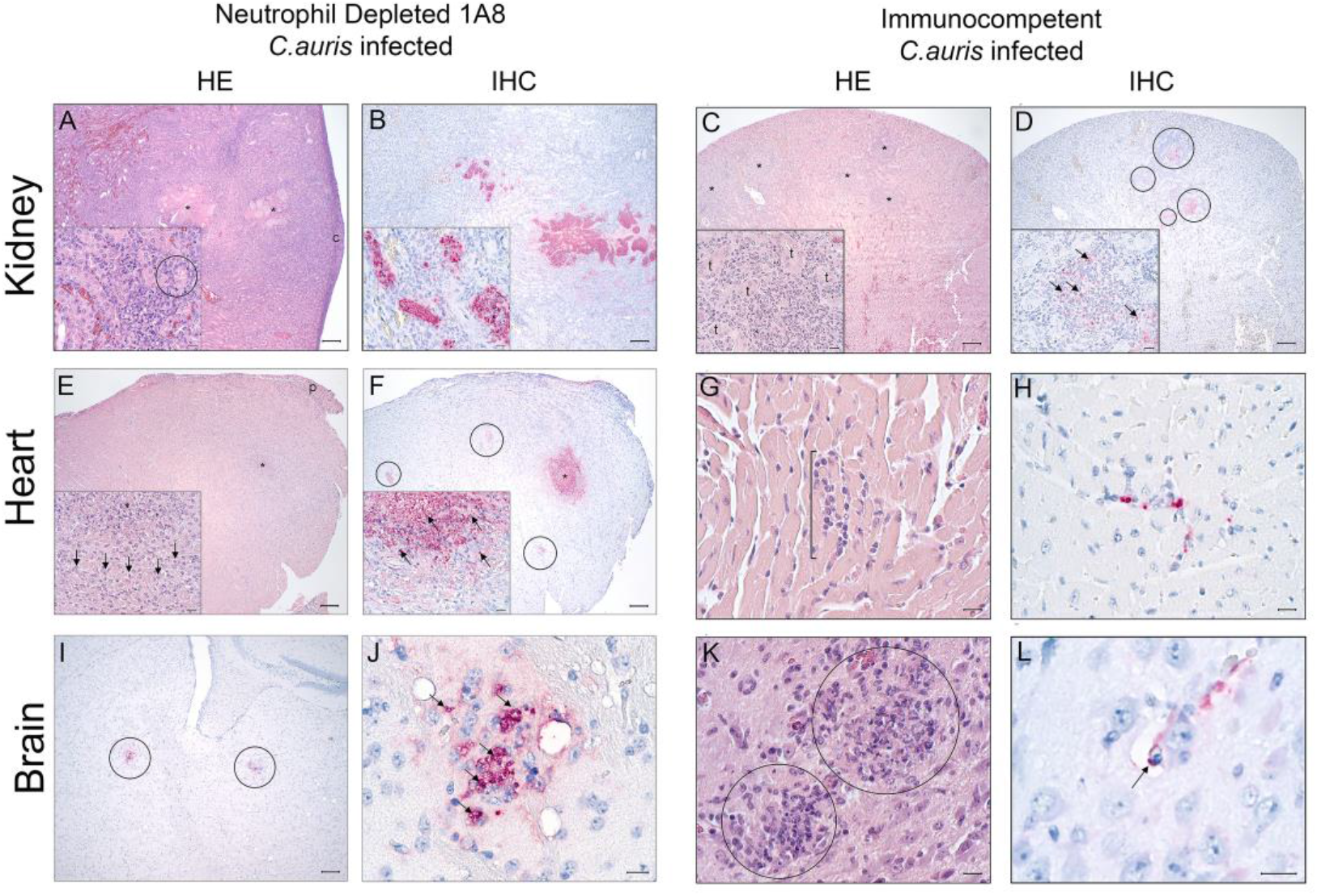
Neutrophil depleted mice infected with reveal microabscesses in target organs, kidney, heart and brain. Histopathology (H&E) and corresponding *C. auris* Immunohistochemistry (IHC) in organs of animals treated with antibody 1A8 neutrophil depleted mice were and inoculated i.v. with 10^7^ *C. auris* cells IHC (B,F,J); positive cells and antigen are highlighted in red. Magnification of images A,B.E,F, I is 5x (bar = 100 μm), inserts 50x (bar = 10 μm); J is 60x (bar = 10 μm). For 1A8 treated mice, Kidney (A-B); Coalescing abscesses with necrotic cores (*) expanding to the renal capsule (c) and abundant intralesional yeast (insert-circle). Prominent antigen staining in areas of inflammation and lumina of renal tubules (insert). Heart (E-F); Large myocardial abscess (*) with extensive diffuse inflammation extending to the pericardium (p). Inflammatory infiltrate consists mostly of macrophages (insert-arrows). Antigen in abscess (*) and microabscesses (circles). Numerous intact yeasts in abscess (insert). Brain (I-J); Microabscesses highlighted by the IHC (circle). Focus with *C. auris* clusters and single yeasts (arrows). For immunocompetent mice, histopathology (H&E) and corresponding *C. auris.* Immunohistochemistry (IHC) in organs of animals inoculated i.v. with 10^7^ units of *C. auris*. IHC (D,H, and L); positive cells and antigen are highlighted in red. Magnification of images C-D is 5x (bar = 100 μm), G-K and inserts are 50x (bar = 10 μm), L is 100x (bar = 10 μm). Kidney (C-D); Foci of inflammation (*) with numerous neutrophils infiltrating the interstitium between renal tubules (insert-t). Prominent antigen staining in areas of inflammation (circles) and intact yeast admixed with suppurative infiltrate (insert-arrows). Heart (G-H); Focus of neutrophils and macrophages infiltrating the myocardium (bracket). Intact yeast between myofibers and surrounded by inflammatory cells. Brain (E-F); Foci of neutrophils, macrophages and microglial (circle). Intact yeast in a capillary perivascular space (arrow).

**Table 1.**
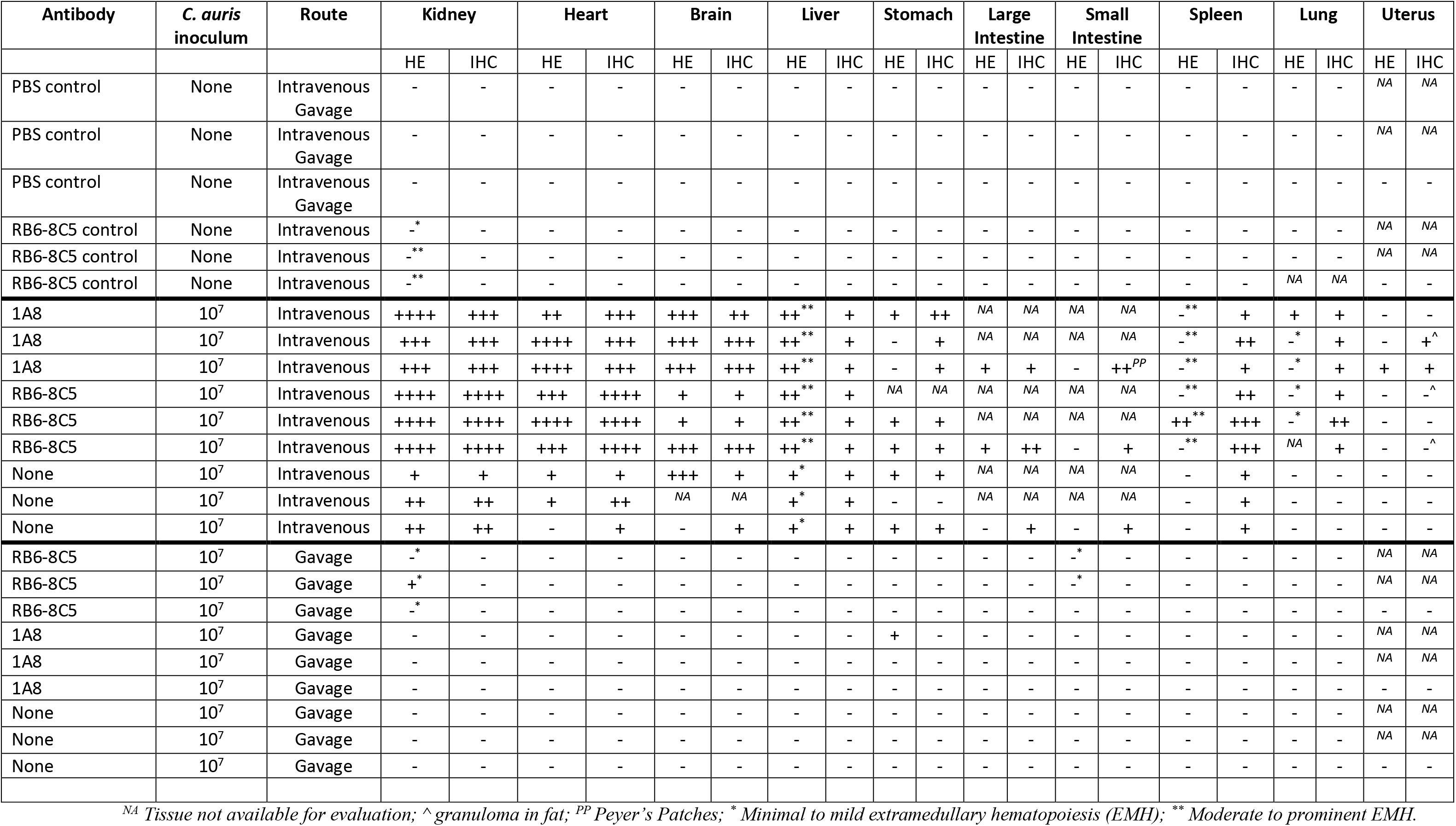
Fungal organ burden scoring system for neutrophil depleted and immunocompetent BALB/c mice infected with *C. auris*. A comparison of route of infection and neutrophil depletion shows the following: **Lesion scoring (HE):** - WNL; + Minimal to mild suppurative inflammation, scattered macrophages and rare microabscessation; ++ Mild to moderate suppurative inflammation, moderate numbers of macrophages and microabscesses; +++ Severe multi-focal necrotizing suppurative inflammation, numerous macrophages microabscesses; ++++ Severed multi-focal-to-coalescing necrotizing suppurative inflammation, numerous macrophages and microabscesses. **IHC:** - negative; + 1-5 cells per high power field—HPF (50x); ++ 5-10 cell/HPF and scattered antigen in debris;+++ 10-20 cells/HPF and scattered antigen in debris; ++++ > 20 cells/HPF and scattered antigen in debris.

| Antibody | *C. auris*<br>inoculum | Route | Kidney |  | Heart |  | Brain |  | Liver |  | Stomach |  | Large Intestine |  | Small Intestine |  | Spleen |  | Lung |  | Uterus |  |
| --- | --- | --- | --- | --- | --- | --- | --- | --- | --- | --- | --- | --- | --- | --- | --- | --- | --- | --- | --- | --- | --- | --- |
|  |  |  | HE | IHC | HE | IHC | HE | IHC | HE | IHC | HE | IHC | HE | IHC | HE | IHC | HE | IHC | HE | IHC | HE | IHC |
| PBS control | None | Intravenous<br>Gavage | - | - | - | - | - | - | - | - | - | - | - | - | - | - | - | - | - | - | NA | NA |
| PBS control | None | Intravenous<br>Gavage | - | - | - | - | - | - | - | - | - | - | - | - | - | - | - | - | - | - | NA | NA |
| PBS control | None | Intravenous<br>Gavage | - | - | - | - | - | - | - | - | - | - | - | - | - | - | - | - | - | - | - | - |
| RB6-8C5 control | None | Intravenous | -* | - | - | - | - | - | - | - | - | - | - | - | - | - | - | - | - | - | NA | NA |
| RB6-8C5 control | None | Intravenous | -** | - | - | - | - | - | - | - | - | - | - | - | - | - | - | - | - | - | NA | NA |
| RB6-8C5 control | None | Intravenous | -** | - | - | - | - | - | - | - | - | - | - | - | - | - | - | - | NA | NA | - | - |
| 1A8 | 10<sup>7</sup> | Intravenous | ++++ | +++ | ++ | +++ | +++ | ++ | ++** | + | + | ++ | NA | NA | NA | NA | -** | + | + | + | - | - |
| 1A8 | 10<sup>7</sup> | Intravenous | +++ | +++ | ++++ | +++ | +++ | +++ | ++** | + | - | + | NA | NA | NA | NA | -** | ++ | -* | + | - | +<sup>^</sup> |
| 1A8 | 10<sup>7</sup> | Intravenous | +++ | +++ | ++++ | +++ | +++ | +++ | ++** | + | - | + | + | + | - | ++<sup>PP</sup> | -** | + | -* | + | + | + |
| RB6-8C5 | 10<sup>7</sup> | Intravenous | ++++ | ++++ | +++ | ++++ | + | + | ++** | + | NA | NA | NA | NA | NA | NA | -** | ++ | -* | + | - | -<sup>^</sup> |
| RB6-8C5 | 10<sup>7</sup> | Intravenous | ++++ | ++++ | ++++ | ++++ | + | + | ++** | + | + | + | NA | NA | NA | NA | ++** | +++ | -* | ++ | - | - |
| RB6-8C5 | 10<sup>7</sup> | Intravenous | ++++ | ++++ | +++ | ++++ | +++ | +++ | ++** | + | + | + | + | ++ | - | + | -** | +++ | NA | + | - | -<sup>^</sup> |
| None | 10<sup>7</sup> | Intravenous | + | + | + | + | +++ | + | + | + | + | + | NA | NA | NA | NA | - | + | - | - | - | - |
| None | 10<sup>7</sup> | Intravenous | ++ | ++ | + | ++ | NA | NA | + | + | - | - | NA | NA | NA | NA | - | + | - | - | - | - |
| None | 10<sup>7</sup> | Intravenous | ++ | ++ | - | + | - | + | + | + | + | + | - | + | - | + | - | + | - | - | - | - |
| RB6-8C5 | 10<sup>7</sup> | Gavage | -* | - | - | - | - | - | - | - | - | - | - | - | -* | - | - | - | - | - | NA | NA |
| RB6-8C5 | 10<sup>7</sup> | Gavage | + | - | - | - | - | - | - | - | - | - | - | - | -* | - | - | - | - | - | NA | NA |
| RB6-8C5 | 10<sup>7</sup> | Gavage | -* | - | - | - | - | - | - | - | - | - | - | - | - | - | - | - | - | - | - | - |
| 1A8 | 10<sup>7</sup> | Gavage | - | - | - | - | - | - | - | - | + | - | - | - | - | - | - | - | - | - | NA | NA |
| 1A8 | 10<sup>7</sup> | Gavage | - | - | - | - | - | - | - | - | - | - | - | - | - | - | - | - | - | - | NA | NA |
| 1A8 | 10<sup>7</sup> | Gavage | - | - | - | - | - | - | - | - | - | - | - | - | - | - | - | - | - | - | - | - |
| None | 10<sup>7</sup> | Gavage | - | - | - | - | - | - | - | - | - | - | - | - | - | - | - | - | - | - | NA | NA |
| None | 10<sup>7</sup> | Gavage | - | - | - | - | - | - | - | - | - | - | - | - | - | - | - | - | - | - | NA | NA |
| None | 10<sup>7</sup> | Gavage | - | - | - | - | - | - | - | - | - | - | - | - | - | - | - | - | - | - | - | - |

<sup>NA</sup> Tissue not available for evaluation; <sup>^</sup> granuloma in fat; <sup>PP</sup> Peyer's Patches; \* Minimal to mild extramedullary hematopoiesis (EMH); \*\* Moderate to prominent EMH.

Damage in the liver ranged from mild in infected control mice to moderate in infected 1A8 and RB6-8C5 treated mice. We observed a diffuse suppurative hepatitis, characterized by moderate numbers of neutrophils in the sinusoids and rare microabscessation in the treated groups. Changes in the gastrointestinal track were observed in all groups and consisted of minimal and multi-focal infiltration of neutrophils and scattered macrophages in the gastric mucosa and submucosa. Similar changes were observed in the large intestine of both treatment groups. Pathology in the spleen was restricted to one animal that was treated with RB6-8C5 and consisted of a large abscess. While the urinary bladder and uterus of all animals were within normal limits, microgranulomas were present in the endometrium and peritoneal adipose tissue associated to the uterus of treated animals (Table 1 and Fig. S3).

We also found various degrees of extramedullary hematopoiesis (EMH) in the liver, spleen, and lungs, which was more prominent in infected 1A8 and RB6-8C5 treated mice. Animals treated with RB6-8C5 and challenged i.v. with PBS or by gavage with *C. auris*, also exhibited various degrees of EMH in the liver and lungs (data not shown). Overall, tissues from control animals challenged i.v. with PBS or with *C. auris* by gavage, regardless of the treatment group, were within normal limits (data not shown).

A primary rabbit polyclonal antibody against *C. albicans* that we found to cross react with *C. auris*, allowed us to analyze the tissues by immunohistochemistry (IHC). Measures used to assure the sensitivity and specificity of the assay included, positive and negative controls in each case, along with the positive correlation of solely yeast form in tissue. Positive immunostaining consisted of discrete red staining of yeasts and granular-to-homogenous intracytoplasmic stain which may be *C. auris* secreted or processed antigen. Mirroring histopathological changes, the most prominent antigen distribution was observed in the kidney, heart, and brain. Overall, antigen was detected in areas of inflammation, cellular debris in abscesses, and intracytoplasmic in infiltrating neutrophils and macrophages. Additionally, numerous renal tubular epithelial cells where IHC-positive and numerous intact yeasts were highlighted in the lumen of renal tubules. Parallel to the histopathology, antigen distribution in the brain was more prominent in animals treated with the 1A8 antibody. In addition to the staining in areas of inflammation and meninges, a discrete 253 cytoplasmic staining was also observed in the perivascular space and adventitia of capillaries, 254 interpreted to be astrocytes’ podocytes (Fig. 4, Fig 5 and Table 1).

**Figure 5.**
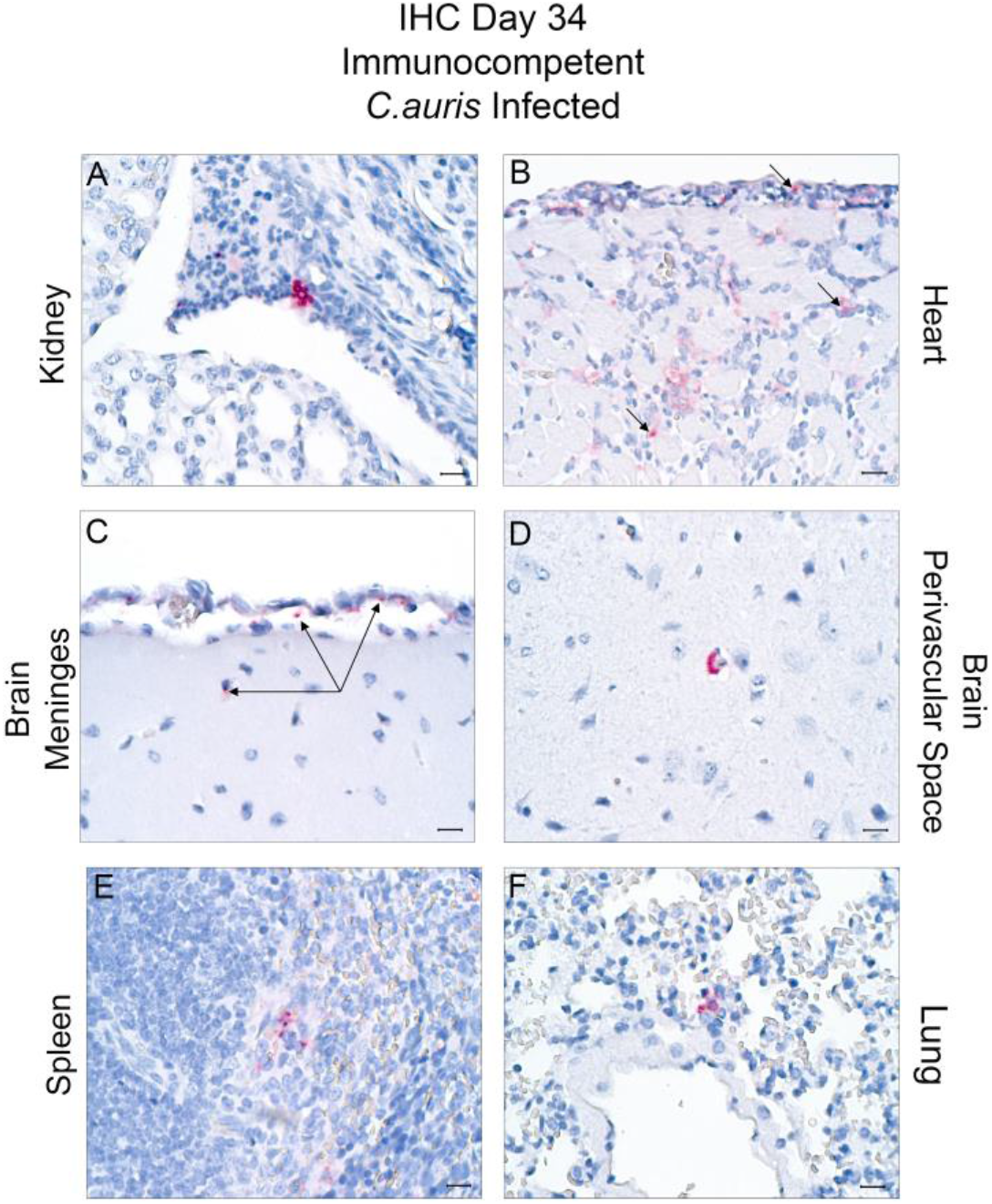
*Candida auris* remains present in immunocompetent mice 34 days post-infection. *C. auris* Immunohistochemistry (IHC) in organs of animals inoculated IV with 10^7^ units of *C. auris* and allowed to clinically recover. Positive cells and antigen are highlighted in red. Magnification is 50x (bar = 100 μm). Kidney (A); Clusters of yeasts in the renal pelvis admixed with inflammatory cells and cell debris. Heart (B); Intact yeast admixed with macrophages between myofibers. Brain (C) Intact yeasts (arrows)in the meninges (D) and perivascular space. Spleen (E); Intact yeast in the red pulp. Lungs (F); Intracytoplasmic antigen in pulmonary macrophage.

Immunostaining was not prominent in liver tissues. We found few scattered neutrophils, macrophages, and microabscesses were positive, and rarely intact yeast in sinusoids. In the gastrointestinal tract, antigen was also highlighted in the Peyer’s patches and lymphoid follicles of the small intestine in all groups (Fig. S3). *C. auris* antigen was also detected in the spleen of all neutrophil depleted and infected control mice but notably more prominent in those treated with RB6-8C5, followed by 1A8 and non-treated consecutively. In the lungs, antigen was only detected in the treated groups (Table 1). Lastly, antigen was detected in the endometrium of 1A8 and RB6-8C5 treated animals as well as in peritoneal fat microabscesses (Table 1 and Fig S3). Additionally, 34 days post infection, in infected control mice, we found the presence of *C. auris* yeast and antigen in kidney, heart, brain, lung, spleen, meninges, stomach, PPs and uterus (Fig. 5).

In humans *C. auris* has been identified to be excreted in urine. Histopathology indicated that the organism was present in the renal tubes and pelvis in our model (1). We tested urine and feces either by qPCR or culture to determine whether mice were shedding *C. auris*. The results reveal that viable *C. auris* is shed in urine and we were able to detect an output of 10^2^ CFU/μl from mice that had initially been infected i.v. with 10^7^ *C. auris* cells (Fig. 6A). We also found *C. auris* in the feces ranging from 10^4^-10^5^ CFU/ μl in both the i.v. and gavage models. We noted that neutrophil depleted mice with RB6-8C5 had a one log difference in CFUs (Fig. 6A). qPCR performed in feces revealed that *C. auris* i.v. infected mice had a cycle threshold value (Ct) of 33.18±1.65 for RB6-8C5. Infection by gavage showed a Ct value 31.7±1.81 for RB6 (Fig. 6). Based on our cycle threshold (Ct) curve analysis, a cycle threshold (Ct) value of ≤38 was reported as positive and ≥38 was reported as a negative. To determine if the fecal microbiota changed upon infection, we performed 16S sequencing that allowed us to interrogate the fecal microbiome. The results revealed that systemic *C. auris* infection does not change the composition of the microbiome (Fig. 7).

**Figure 6.**
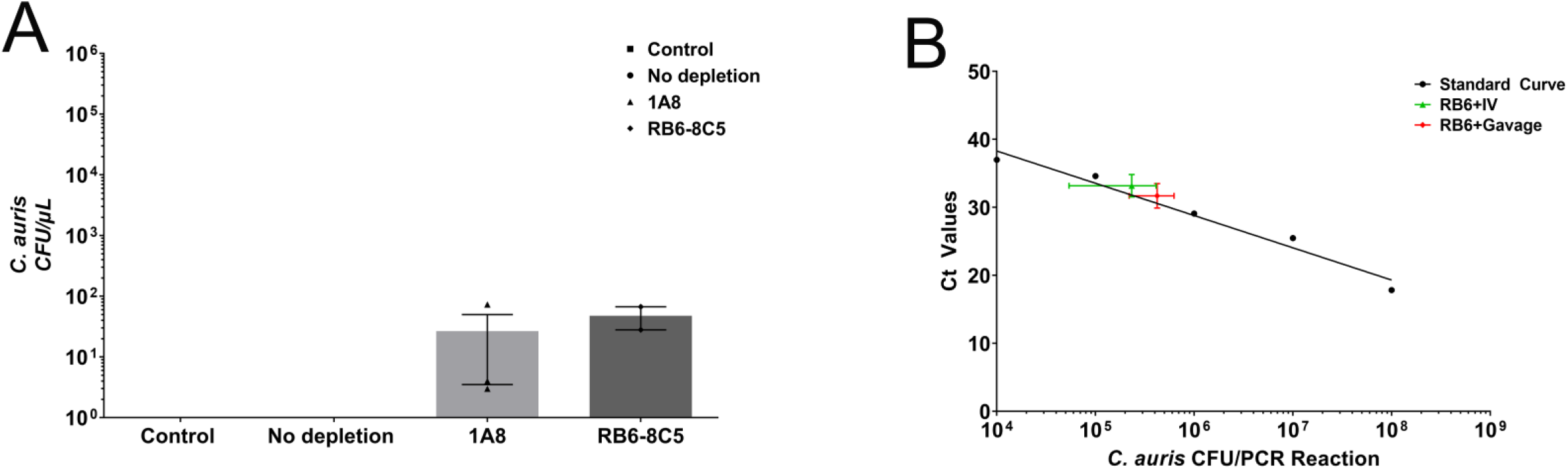
*C. auris* is found in urine and feces 7 days post-infection. (A)Neutrophil depleted mice with either 1A8 or RB6-8C5 infected with 10^7^ *C. auris* by i.v. were found positive by culture on sabouraud dextrose agar plates. The *C. auris* output in urine was found to be 10^2^ CFU/μL. (B) By qPCR fecal pellets from RB6-8C5 neutrophil depleted mice infected with 10^7^ *C. auris* by i.v. show an output 10^5^ the gavaged mice show an output of 10^6^.

**Fig. 7.**
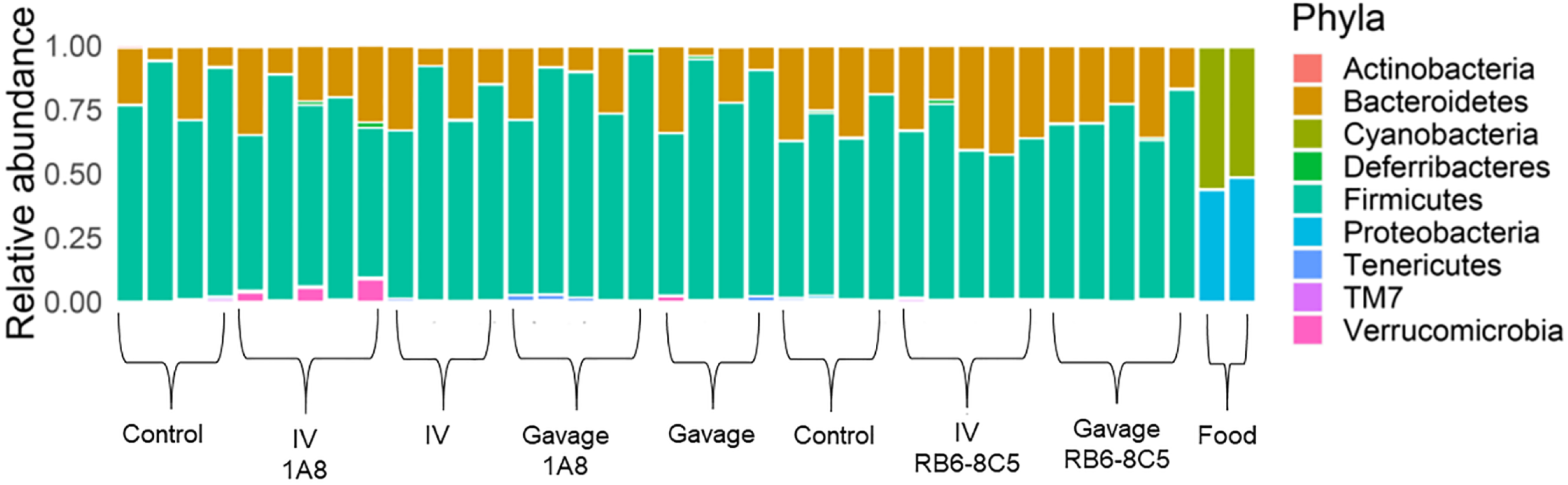
The fecal microbiota is not altered neutrophil depleted mice. Fecal DNA was extracted and Illumina 16S next generation sequencing was performed. Regardless of which antibody 1A8 or RB6-8C5 used to depleted neutrophils the fecal microbiota remained consistent. The route of infection also did not alter the microbial composition.

Regardless of whether mice were neutrophil depleted or the infected control, they developed a head tilt known as torticollis as previously observed at 8-13 post i.v. infection with *C. auris* during our survival experiments (21). We observed that between days 8-13, 50% of the mice exhibiting torticollis oriented to the right and 50% oriented to the left (Fig. 8 and Movie 1). When these mice were elevated by the tail they exhibited a forceful spin on their tails until they were set back down (Fig. 8 and Movie 2). Twenty-two days after the torticollis appears, 50% of the mice exhibited a rapid head bobbing along with fast and erratic movement in the cage (Fig. 8 and Movie 3). We found that a head bobbing does not revert back to torticollis. When these mice are then elevated by their tails the head bobbing mice exhibit a curling like behavior (Fig.8 and Movie 4). Interestingly, the head bobbing behavior does not revert to torticollis. Histology of the cerebellum an area important in balance, coordination and muscular activity did not show the presence of *C. auris* therefore further analysis of the inner ear is required to understand this phenotype. Surviving control mice that were infected with 10^7^ and 10^8^ *C. auris* cells were followed for 104 days post infection. These mice continued to exhibit the head bobbing along with fast and erratic movement in the cage (Movie 5).

**Figure 8:**
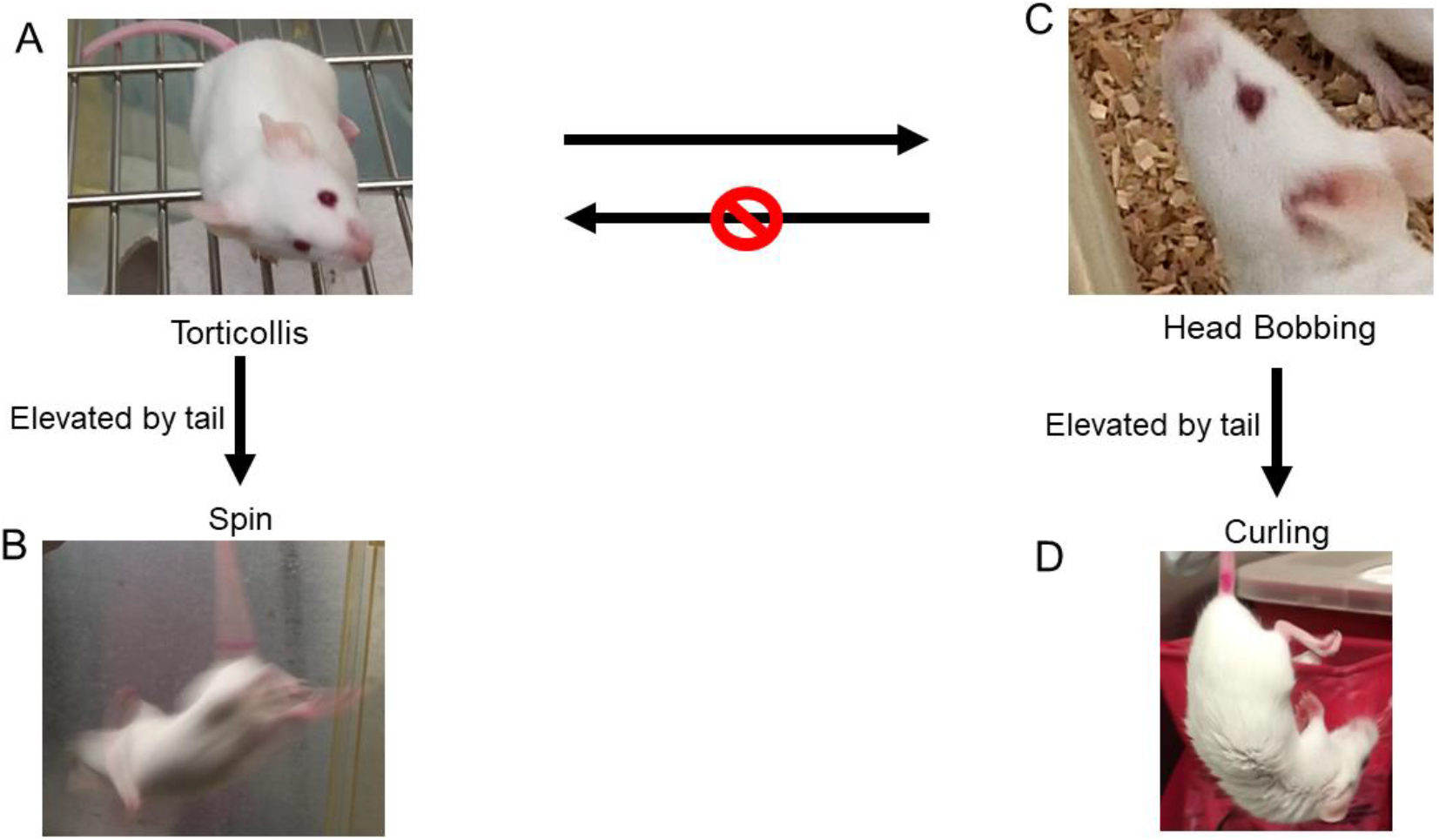
Progression of behavioral phenotype. Mice infected with 10^7^ *C. auris*, regardless of whether they were immunocompetent or neutropenic, displayed a behavioral phenotype. 8-13 days post infection mice displayed torticollis (characterized by a lean) when these mice were elevated by the tail they began to spin. By day 22 post infection, mice displayed head bobbing movement, these mice when elevated by the tail showed curling. Mice exhibiting f head bobbing did not appear to revert by to torticollis.

## Discussion

Since its identification in 2009, *Candida auris* has become a major public health threat due to its unpredictable multidrug resistance profile, high transmissibility, particularly in nursing home and healthcare settings and high mortality rates of 30%-60%. Currently, there are a limited number of murine models of *C. auris* infection that can be used to study disease progression. Although there are immunocompetent models of infection there are discrepancies in survival based on the murine strains and isolates used (21, 22). The immunosuppressed models use cyclophosphamide, which depletes the entire immune system to allow for *C. auris* dissemination but limits studies that test host immunity against this pathogen(24). Here we compare two mouse models that achieve immunosuppression by depleting neutrophils using 1A8, anti-Ly6G^+^ which depletes neutrophils only and RB6-8C5, anti- Ly6G^+^-Ly6C^+^ which depletes neutrophils and Ly6G monocytes(30). We chose to deplete neutrophils as it was recently documented that neutrophils poorly recruit *C. auris* and failed to form neutrophil extracellular traps (NETs), structures of DNA, histones, and proteins with antimicrobial activity (32). Since *C. auris* evades neutrophil attack, the 1A8 and RB6-8C5 models can be used to further elucidate how *C. auris* takes advantage of the host in the absence of neutrophils (32). Both models were useful as these as 1A8 represents long term depletion while RB6-8C5 represents short term depletion.

Our models also confirm that an inoculum of 10^7^ *C. auris* cells is optimal to study *C. auris* progression as previously documented (21, 22). We found that the major target organs were kidney, heart and brain which recapitulates human infection where patients have been found to have kidney failure and endocarditis (14–17). The presence of *C. auris* in kidney tubules detected by histology as well as the presence in urine and feces shows that the neutrophil depletion models recapitulate shedding in humans (33). By histology we found that in addition to yeast cells in tissues, there was the presence of granular-to-homogenous intracytoplasmic stain which may be secreted by *C. auris* or processed antigen; an area to be further investigated. Interestingly, we noted that between 8-13 days post *C. auris* infection, mice displayed a unique behavioral phenotype characterize by torticollis and tail spinning when elevated and that this phenotype can progress to head bobbing, erratic fast movements and body curling by day 22 post infection. While these changes have not been reported in humans, two clinical studies described patients admitted with altered mental status and unsteady gait (14, 18). However, there was no clear indication whether this was due to *C. auris* infection and this is an area where further studies are needed (14, 18). Although *C. auris* microabsesses were present in the brain, it was not present in the cerebellum an area important in balance and coordination therefore, further studies will need to be done to address the pathology of the inner ear. The neutrophil depletion models allude to whether mice are dying from *C. auris* or with *C. auris* and this depends on the state of the host. The neutrophil depletion model suggest that the host dies from *C. auris*, the immune competent infected control model suggests that *C. auris* stays present in the host without causing disease and may become opportunistic upon a change in immune status. Further studies are required to understand how this fascinating organism detects the immune status of the host.

## Materials and Methods

### Animal Use Ethics Statement

All animal experiments in this study were reviewed and approved by the Wadsworth Center’s Institutional Animal Care and Use Committee (IACUC) under protocol #18-450 and is fully accredited by the Association for Assessment and Accreditation of Laboratory Animal Care (AAALAC). The Wadsworth Center complies with the Public Health Service Policy on Humane Care and Use of Laboratory Animals and was issued assurance number A3183-01.

### Animals

BALB/c female mice (8-12 weeks old) were obtained from Taconic Farms (Hudson, NY). Animals were housed in autoclaved micro-isolator cages under specific pathogen-free conditions and were treated in compliance with the Wadsworth Center’s Institutional Animal Care and Use Committee (IACUC) guidelines. The BALB/c mice were housed in an animal holding room and all procedures were conducted in a biosafety cabinet. Standard operating procedures were followed as recently described in (12).

### Neutrophil Depletion and Infection with *C. auris*

Neutropenia was induced in mice by intraperitoneal (i.p.) injection of 200 μg of monoclonal antibodies 1A8 (Bio X Cell, West Lebanon, NH) that targets Ly6G cells or RB6-8C5(Bio X Cell, West Lebanon, NH) that targets Ly6C and Ly6G cells. Injections with the antibodies were administered 24 hours prior to infection and every 48 hours thereafter. Tail vein and gavage infections were done with10^7^ or 10^8^ *C. auris* strain CAU-09 acquired from the available Center’s for Disease Control (CDC) panel. For survival studies, infected mice were weighed daily, and the development of illness was monitored. Mice were euthanized when they demonstrated signs of morbidity or lost 20-25% of their body weight.

### Fecal pellet and urine collection

Fecal pellets and urine were collected before initial neutrophil depletion, before infection, and prior to euthanasia.180-220 mg (4-5 fecal pellets) were collected for DNA isolation, snap frozen in liquid nitrogen and stored at 80°C. 20-100 μl of urine was collected and 1μl was immediately plated onto Sabouraud dextrose agar plate for with antibiotics (Chloramphenicol (20 μg/ml), Penicillin (20U), Gentamycin (40 μg/ml), Streptomycin (40 μg/ml) for 24 h at 38°C to determine CFU’s, the remainder was stored at −20°C.

### Tissue Collection for CFU counts and Histology

Mice were euthanized by CO_2_ and cardiac punctures were performed. Blood was centrifuged at 16,000 rpm for 2 min and serum was stored at −20°C. Necropsies of the bladder, uterus, spleen, kidney, cecum, small intestine, stomach, liver, heart, lung and brain were collected and weighed for each mouse and used for CFU counts and histology. For CFU counts, the organs were homogenized in Hanks Sodium Balanced Salt (HBSS) (Thermo Fisher, Waltham, MA) through a 70μm cell strainer (Cell Treat, Pepperell, MA). A 1:100 dilution of organ homogenate was plated onto Sabouraud dextrose agar plate with antibiotics (Chloramphenicol (20μg/ml), Penicillin (20U), Gentamycin (40μg/ml), Streptomycin (40μg/ml)). CFU/gram of each organ was determined based on *C. auris* growth. For histology, organs were placed in 10% formalin for 24 hours followed by 70% ethanol before being submitted to the histology core facility for processing.

### Histopathology

A representative sample of liver, kidney, heart, brain, stomach, spleen, lung, uterus, urinary bladder, small and large intestines were collected during necropsy and fixed in 10% buffered formalin for 24 hours. Tissues were subsequently transferred to 70% ethanol prior to processing and embedding. Tissue sections were stained with hematoxylin and eosin for histopathological evaluation.

### Immunohistochemistry (IHC)

Tissue sections, 3-4μm in thickness, were deparaffinized in CitriSolve (Decon Labs., King of Prussia, PA) and rehydrated by processing them through graded alcohols. Tissues were pretreated with Proteinase K (10 μg/mL) for 15 minutes. Endogenous IgG and non-specific background were blocked with Rodent Block M (Biocare Medical; Pacheco, CA) for 20 minutes, followed by an alkaline phosphatase block (BLOXALL; Vector Laboratories, Burlingame, CA) for 10 minutes. The primary antibody (PA1-7206; Thermo Fisher Scientific, Waltham, MA) was incubated on the tissue sections at a dilution of 1: 10,000 for 1 hour at room temperature. Subsequently, sections were sequentially incubated with a rabbit-on-rodent tissue alkaline phosphatase-based polymer and Warp Red (Biocare Medical, Pacheco, CA)). Tissues were counterstained with Tacha’s hematoxylin and mounted using EcoMount (Biocare Medical, Pacheco, CA).

### Fecal DNA purification

DNA from the fecal samples was isolated using Qiagen AllPrep® PowerFecal® DNA/RNA Kit according to manufacturer’s instructions. The only minor modification that was made was that fecal samples were lysed with provided lysis buffer using a vertical Disrupter Genie for 5 minutes at 3000 rpm. Fecal DNA samples were measured by Qubit quantification and aliquots were used for qPCR and sequencing. The remainder of the DNA samples were stored at −20°C.

### ITS2 Fungal qPCR

The fecal DNA sample were subjected qPCR. The reactions were done using *ITS2* primer for fungal genomes (34). This PCR confirms that the presence of *C. auris* fecal DNA. A standard curve was produced by performing a serial dilution of 10^8^ to 10^−1^ *C. auris* cells. These dilutions were then processed identically to the fecal samples collected to extract and purify DNA and RNA. A TaqMan qPCR reaction was run on the DNA samples using the *ITS2* primer and probes for *C. auris*. The PerfeCTa Multiplex qPCR ToughMix was spiked with bicoid plasmid and respective primers after pipetting of the negative template control wells. Each sample and control were run in duplicate on a 7500 Fast instrument for 45 cycles. A standard curve was generated and used to calculate relative concentrations of *Candida auris* in the fecal pellets depending on their cycle threshold value. The fecal pellet DNA samples were run identically to the standard curve samples.

### 16S Sequencing and analysis

Ilumina 16S metagenomic sequencing library was prepared as described in manufacturer’s instructions. The16S amplicon forward and reverse primers from Illumina were used in first round of PCR for 8 cycles. The samples were cleaned using AMPure XP beads and second round of index PCR was performed for an additional 8 cycles. The samples were cleaned, quantified, pooled and subjected to sequencing using Illumina MiSeq system. The analysis of the sequencing data was performed as follows: BBMerge from BBMap v.37.93 (https://sourceforge.net/projects/bbmap/) simultaneously quality trimmed and merged paired-end reads under the following parameters: reads were required to be a minimum of 100 or 150 base pairs in length with an average PHRED quality score of 20 and a quality cutoff of 20 per position, merged reads were discarded if the overlap between them contained three times the expected number of errors. BBDuk from BBMap removed forward and reverse primers from 16S sequences using a k-mer size of 15 and a minimum k-mer size of 11. Seqtk v.1.3 (https://github.com/lh3/seqtk) converted fastq files to a fasta format and all sequences were merged into a single multi-fasta file for OTU assignment and taxonomic classification in NINJA-OPS v.1.5.1(35). Briefly, Bowtie2 v.2.1.0 (36) aligned 16S sequences to concatenated 16S Greengenes (37) using the parameters outlined by the NINJA-OPS manual (https://github.com/GabeAl/NINJA-OPS). NINJA-OPS then binned sequences into OTUs and assigned taxonomic information based on sequence alignments to the concatenated databases. Absolute abundances were summarized to the genus level in QIIME v.1.9.0 with the summarize_taxa.py script (38) and significant changes in taxon abundance were calculated in DESEq2 (39) in R v.3.5.2 (http://www.R-project.org).

### Statistical analysis

All summary data are expressed as mean ± SEM. When comparing means from two or more treatment groups, a nonparametric one-way analysis of variance (ANOVA) with Kruskal-Wallis test followed by uncorrected Dunn’s multiple comparison test was done using Prism 6 (GraphPad La Jolla, CA). Statistical significance is indicated in figures using the following denotation: P > 0.05, *P ≤ 0.05, **P ≤ 0.01, ***P ≤ 0.001, ****P ≤ 0.0001.

## Supporting information

Movie 1. Torticollis

Movie 2. Spinning

Movie 3. Head Bobbing Day 22

Movie 4. Curling

Movie 5. Head Bobbing Day 104

## Acknowledgements

We thank Dr. Stuart M. Levitz and Dr. Charles Specht, University of Massachusetts Medical School for their helpful discussions on neutrophil depletion. We thank Yan Zhu and Dr. Sudha Chaurvedi of the clinical mycology laboratory, Wadsworth Center for their help with MALDI-TOF identification of our samples. We thank Dr. Valerie Bolivar, Wadsworth Center for help with statistical analysis for organ fungal burdens. We thank Matthew Shudt and Zhen Zhang at the Wadsworth Center Applied Genomic Technologies Core for technical help of the 16S studies. We thank the Wadsworth Center Histopathology core facility. We also thank Deirdre Torrisi, Frank Blaisdell, Robert Keefe, and the animal care staff at the Wadsworth Center’s Veterinary Sciences Facility for their suggestions and special accommodations that allowed this work to be possible.

## Author contributions

S.R.T., A.P., N.S. and M.D.J. designed and performed the experiments, analyzed and interpreted the data, and wrote the manuscript. F.T.V. stained, analyzed and scored all histopathology sections. R.S. designed and performed the flow cytometry experiments as well as analyzed and interpreted the data. E.L.N. analyzed and graphed the 16S data. M.D.J. supervised the project, oversaw data analyses and interpretation, and wrote the manuscript.

## Data and materials availability

All data is available in the main text or the supplementary materials.

---

**Supplementary Figure 1.**
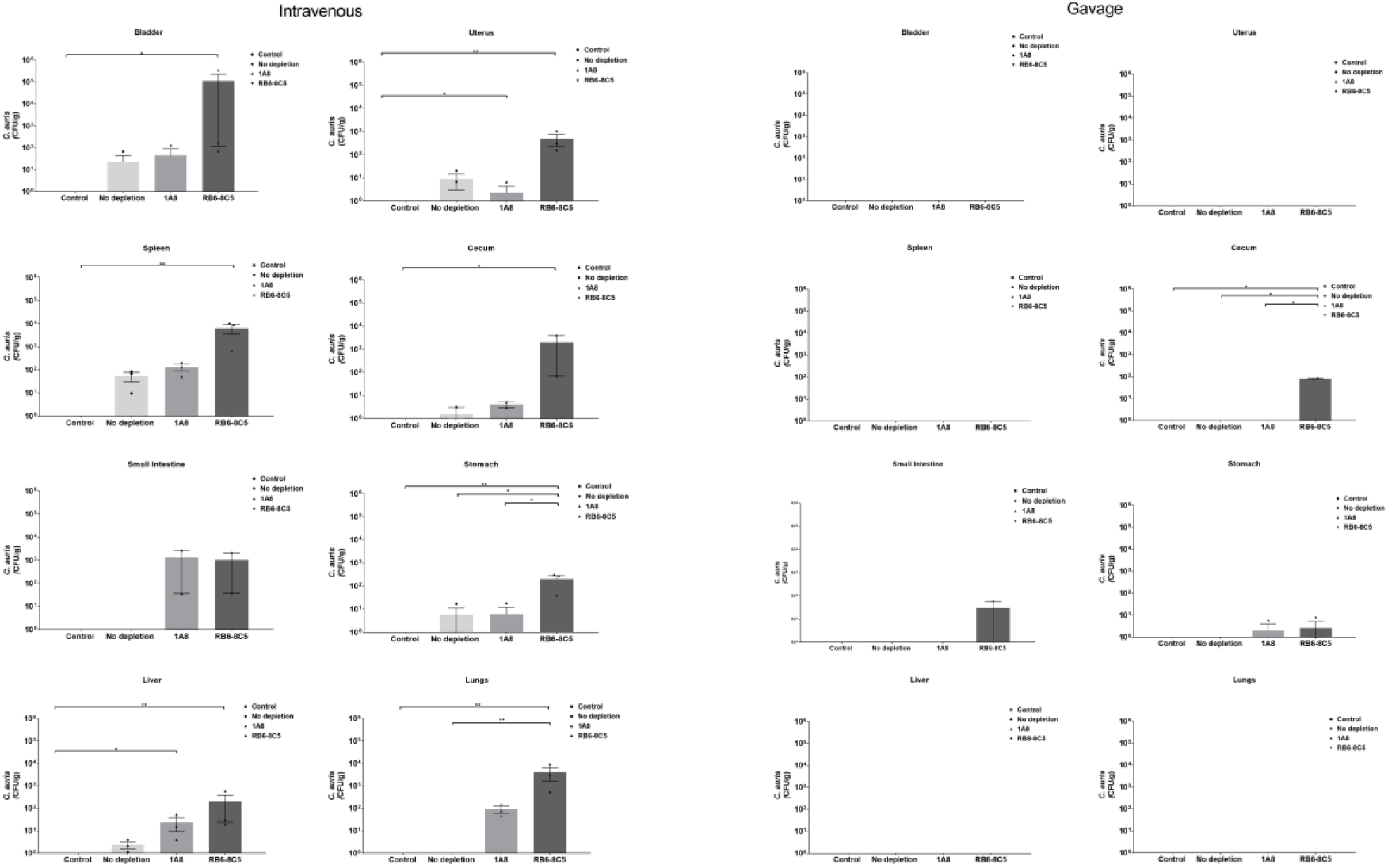
CFU/g of other infected tissues 7 days post-infection. Mice were infected i.v. or by gavage with 10^7^ *C. auris*. Neutropenic mice with 1A8 or RB6-8C5 in the i.v. model showed moderate CFU/g in other organs such as the bladder, uterus,spleen, cecum, small intestine, stomach liver and lungs. Gavaged mice showed modest CFU/g in small intestine and cecum. N=5 mice per group. A nonparametric one-way analysis of variance (ANOVA) with Kruskal-Wallis test followed by uncorrected Dunn’s multiple comparison test was done using ns P > 0.05, *P ≤ 0.05, **P ≤ 0.01, ***P ≤ 0.001, ****P ≤ 0.0001

**Supplementary Fig 2:**
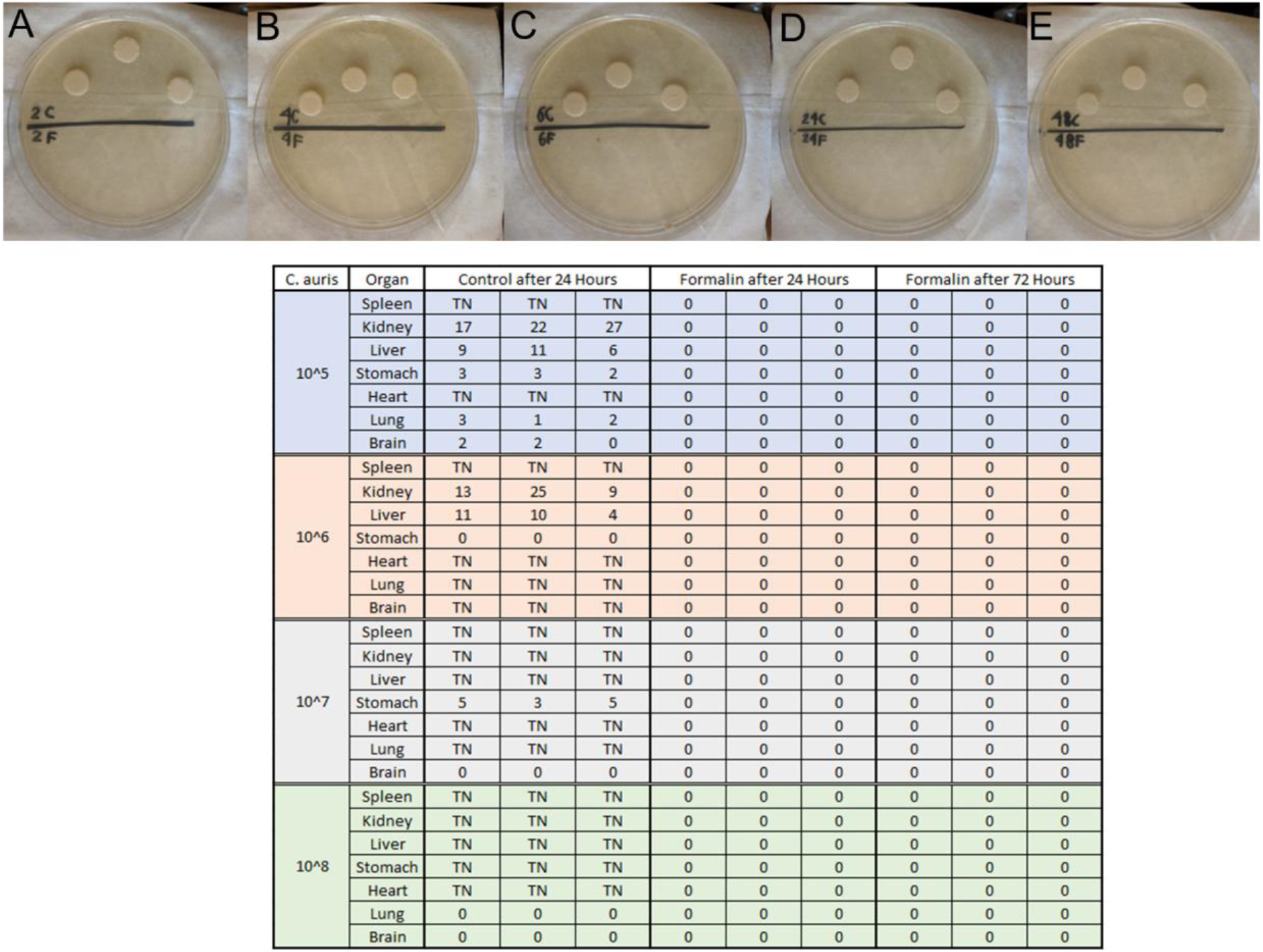
10% Formalin is effective at inactivating *C. auris* in mouse tissues after 24 hours. *C. auris* was incubated in either Sabouraud broth or 10% Formalin at 2, 4, 6, 24, or 48 hours. Within 2 hours, *C. auris* is no longer growing when plated. The table shows tissues collected from mice exposed to different inoculums of *C. auris*. These were incubated in either HBSS or 10% formalin. After either 24 and 72 hours, tissues were washed and plated for growth. There was no growth in tissues incubated for 24 hours in 10% formalin. TN= Too numerous to count.

**Supplementary Figure 3:**
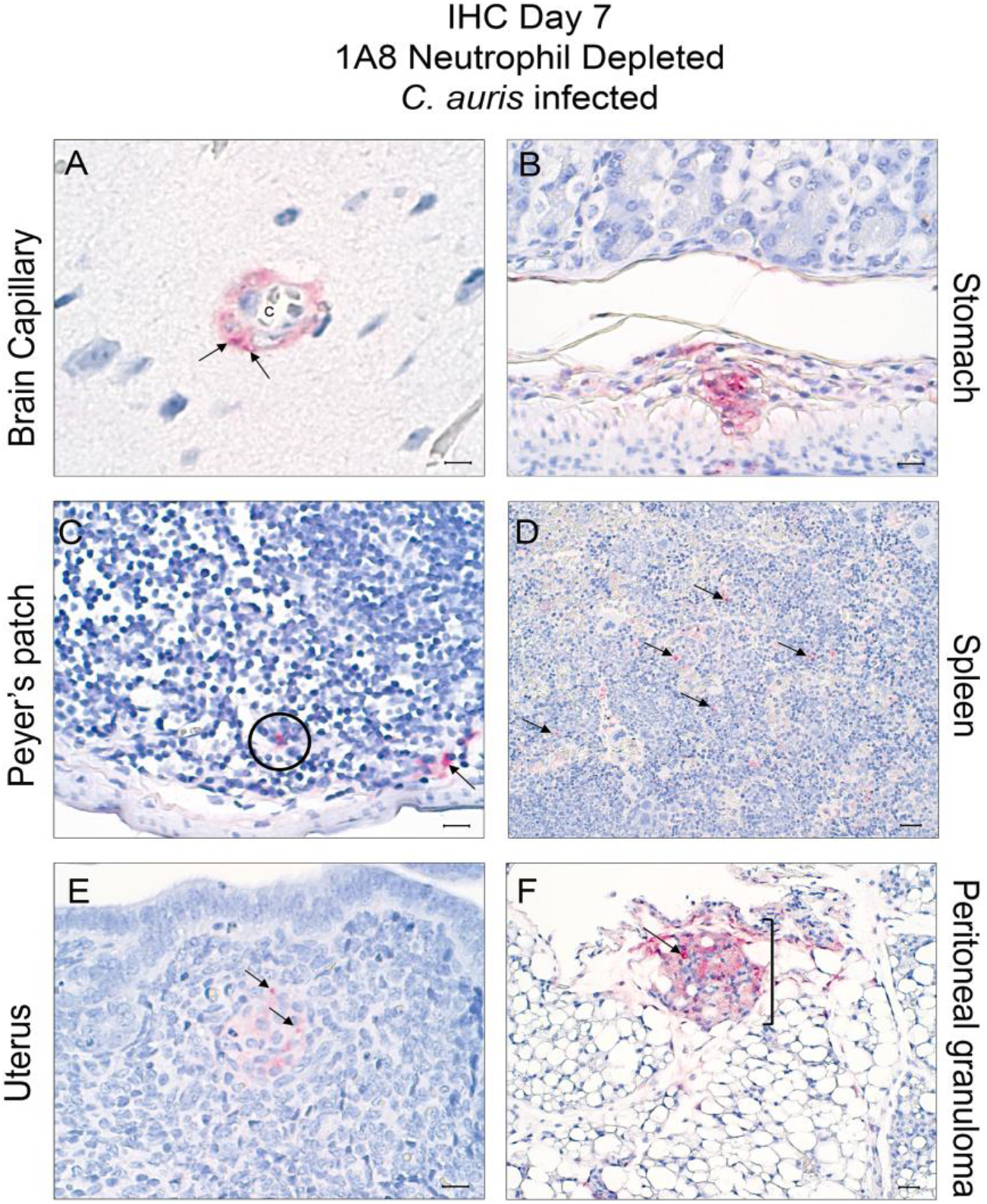
Immunohistochemistry (IHC) of other tissues infected with *C. auris* in 1A8 neutrophil depleted mice, 7 days post-infection. (A) Perivascular cells with intracellular yeast (arrows) in brain capillary of mice (100x / bar 5 μm). (B) Positive staining (red) in the gastric submucosa (50x / bar 10 μm). (C) In the small intestine Peyer’s patches, intracytoplasmic (arrow) and yeast (circle) staining (50x / bar 10 μm). (D) Yeast (arrows) staining in the spleen red pulp in an animal treated with 1A8, (20x / bar 20 μm). (E) Uterine Endometrial granuloma with intralesional yeast (arrows), (50x / bar 10 μm). Peritoneal fat granuloma with intralesional yeast (arrows) (50x / bar 10 μm).

**Supplementary Table 1.** MALDI-TOF confirms the presence of *C. auris* in specific tissues. **Colony forming units from** infected tissues were analyzed by MALDI-TOF to confirm that colonies were *C. auris*. PRC is the positive reaction control. NRC the negative reaction control. CC 70% FA is the color control 70% Formic Acid. PSC is the positive sample control. All tissue samples run were found to contain *C. auris*.

| Sample ID | Organism (Best Match) | Mean |
| --- | --- | --- |
| PRC BTS | *Escherichia coli* | 2.27±0.08 |
| NRC HCCA | **No peaks found** | 0.00 |
| CC 70% FA | **No peaks found** | 0.00 |
| PSC ATCC6258 | *Candida krusei* | 2.4±0.04 |
| 10<sup>7</sup>+RB6-8C5<br>**Heart** | *Candida auris* | 2.39±0.27 |
| 10<sup>7</sup>+RB6-8C5<br>**Lung** | *Candida auris* | 2.375±.028 |
| 10<sup>7</sup>+RB6-8C5<br>**Brain** | *Candida auris* | 2.6±0.04 |
| 10<sup>7</sup>+RB6-8C5<br>**Stomach** | *Candida auris* | 2.52±0.08 |
| 10<sup>7</sup>+RB6-8C5<br>**Kidney** | *Candida auris* | 2.33±0.21 |
| 10<sup>7</sup>+RB6-8C5<br>**Liver** | *Candida auris* | 2.47±0.20 |
| 10<sup>7</sup>+RB6-8C5<br>**Spleen** | *Candida auris* | 2.265±0.32 |

